# Transcriptional terminators allow leak-free chromosomal integration of genetic constructs in cyanobacteria

**DOI:** 10.1101/689281

**Authors:** Ciarán L. Kelly, George M. Taylor, Aiste Satkute, Linda Dekker, John T. Heap

## Abstract

Cyanobacteria are promising candidates for sustainable bioproduction of chemicals from sunlight and carbon dioxide. However, the genetics and metabolism of cyanobacteria are less well understood than model heterotrophic organisms, and the suite of well characterised cyanobacterial genetic tools and parts is less mature and complete. Transcriptional terminators use specific RNA structures to halt transcription and are routinely used in both natural and recombinant contexts to achieve independent control of gene expression and ‘insulate’ genes and operons from one another. Insulating gene expression can be particularly important when heterologous/synthetic genetic constructs are inserted at genomic locations where transcriptional read-through from chromosomal promoters occurs, resulting in poor control of expression of the introduced genes. To date, few terminators have been described and characterised in cyanobacteria. In this work, nineteen heterologous, synthetic or putative native Rho-independent (intrinsic) terminators were tested in the model freshwater cyanobacterium, *Synechocystis* sp. PCC 6803, from which eleven strong terminators were identified. A subset of these strong terminators was then used to successfully insulate a chromosomally-integrated rhamnose-inducible *rhaBAD* expression system from hypothesised ‘read-through’ from a neighbouring chromosomal promoter, resulting in greatly improved inducible control. The addition of validated strong terminators to the cyanobacterial toolkit will allow improved independent control of introduced genes.

## Introduction

Cyanobacteria are important photosynthetic model organisms and potential photosynthetic microbial factories. The ability to reliably engineer photosynthetic organisms could enable the sustainable, carbon-neutral, light-driven conversion of carbon dioxide to valuable products using energy from sunlight. Predictable engineering of cyanobacteria remains challenging for many reasons, one of which is a shortage of well-characterised genetic parts such as promoters, ribosome-binding sites and transcriptional terminators [1,2].

We recently described the successful introduction of a rhamnose-inducible *rhaBAD* expression system into the cyanobacterium *Synechocystis* sp. PCC 6803 (*Synechocystis* hereafter). This expression system has superior properties to many other previously-reported inducible promoter systems, including low basal expression in the absence of inducer, a photostable and non-toxic inducer and a linear response to inducer concentration [3]. However, when introduced into the *Synechocystis* chromosome adjacent to the *ndhB* gene, we observed a non-zero basal level of expression, which we attributed to transcriptional read-through from promoter(s) neighbouring the integration site used. Chromosomal integration is important for the stability of expression constructs, but in cases where extremely low or zero basal expression is required, transcriptional read-through can result in unpredictable gene expression, growth defects, toxicity and genetic instability.

Transcriptional terminators can be used to achieve independent control of gene expression and isolate or ‘insulate’ genes and operons from one another and from neighbouring elements when chromosomally integrated. In contrast to *Escherichia coli*, in which the use and characterisation of transcriptional terminators is routine, evaluation of the function of terminators in *Synechocystis* has been largely ignored to date. There are two main types of terminators: Rho-dependent terminators and Rho-independent or ‘intrinsic’ terminators. Rho-dependent termination requires a homohexameric Rho protein that unwinds the RNA-DNA hybrid, thus halting elongation of nascent RNA strands. No homologues of Rho have been identified in cyanobacterial genomes to date [4]. Rho-independent termination results from the formation of a hairpin-loop secondary structure in the nascent RNA strand, causing dissociation of the transcription elongation complex (comprising RNA polymerase, double-stranded DNA and nascent RNA). Termination is intrinsic to the nucleotide sequence of the RNA strand itself and composed of an adenosine-rich tract (A-tract) located upstream of a hairpin loop consisting of a GC-rich stem region (4-18 bp) and loop nucleotides (3-5 bp), followed by a highly conserved uracil-rich tract (U-tract; 6-8 bp). Transcription of the U-tract transiently pauses the elongation complex, allowing formation of the hairpin loop. This destabilises the complex, resulting in DNA:RNA hybrid shearing, termination of elongation and release of the partial transcript. Interestingly, an analysis of RNA-folding energetics near stop codons in *Synechocystis* suggested a lack of RNA hairpin-loop formation at these sites, implying Rho-independent termination is not prevalent in this cyanobacterium [4,5]. A comprehensive analysis of terminators in *Synechocystis* has not been carried out to date.

In this work we screened nineteen Rho-independent, transcriptional terminator sequences in *Synechocystis*. A subset of the best performing terminators were then tested for their ability to insulate a previously described [3] rhamnose-inducible yellow fluorescent protein (YFP) reporter cassette from likely transcriptional read-through at the site of chromosomal integration. Introduction of any of the chosen terminators resulted in a basal level of YFP production that was indistinguishable from *Synechocystis* cells lacking the YFP expression cassette entirely, confirming successful insulation of chromosomally-integrated constructs.

## Results

### Screening Rho-independent terminators

Rho-independent terminators were identified from the literature including twelve strong, natural terminators from *E. coli* [6] (two of which were taken from the Biobricks registry of biological parts and used previously in cyanobacteria but not tested for efficacy [2,7,8]), four synthetic terminators that showed excellent transcriptional termination in *E. coli* [6], one putative cyanobacterial terminator [9] and T21 and M13 bacteriophage terminators [10,11] (Table S1).

In order to screen them for transcriptional termination activity in *Synechocystis*, the nineteen terminators were inserted immediately downstream of the transcriptional start site (TSS+1) of the *E. coli rhaBAD* promoter and upstream of the ribosome-binding site (RBS) in front of the YFP-encoding gene on plasmid pCK306 [3] (Figure 1A). This *Synechocystis*-*E. coli* shuttle plasmid both replicates stably in *E. coli* and integrates by homologous recombination into the *Synechocystis* genome at a site adjacent to the *ndhB* gene. The *rhaBAD* promoter allows a linear induction of YFP production with respect to the concentration of the inducer (L-rhamnose) added to the growth medium [3]. The resulting plasmids pAS001, pAS002, pAS004-pAS020 were verified by sequencing and initially tested in two commonly-used *E. coli* strains, DH10□ and MG1655. *E. coli t*ransformants were grown in rich media with or without 0.6 mg/ml L-rhamnose at 37 °C and fluorescence was measured after 6 h by flow cytometry, as previously described [3]. YFP production was abolished in all transformants containing the integrated terminator constructs compared to the positive control pCK306 transformants (highly statistically-significant different; *P* value < 0.0001) (Figure S1). No statistically-significant difference in fluorescence was observed between pAS001, pAS002, and pAS004-pAS020 transformants grown with or without inducer. Indeed no statistically-significant difference could be observed between the basal fluorescence of pAS001, pAS002, and pAS004-pAS020 transformants in the presence of inducer and negative control pCK324 transformants, which lack the *rhaBAD* promoter entirely. Subtle differences were observed between the two *E. coli* strains, for example statistically-significant YFP production with inducer was observed with plasmids pAS017 and pAS020 in MG1655 but not in DH10β (P values of < 0.0001 and <0.05 respectively). No changes in these observations were identified after a further 18 h of growth (data not shown).

**Figure 1.**
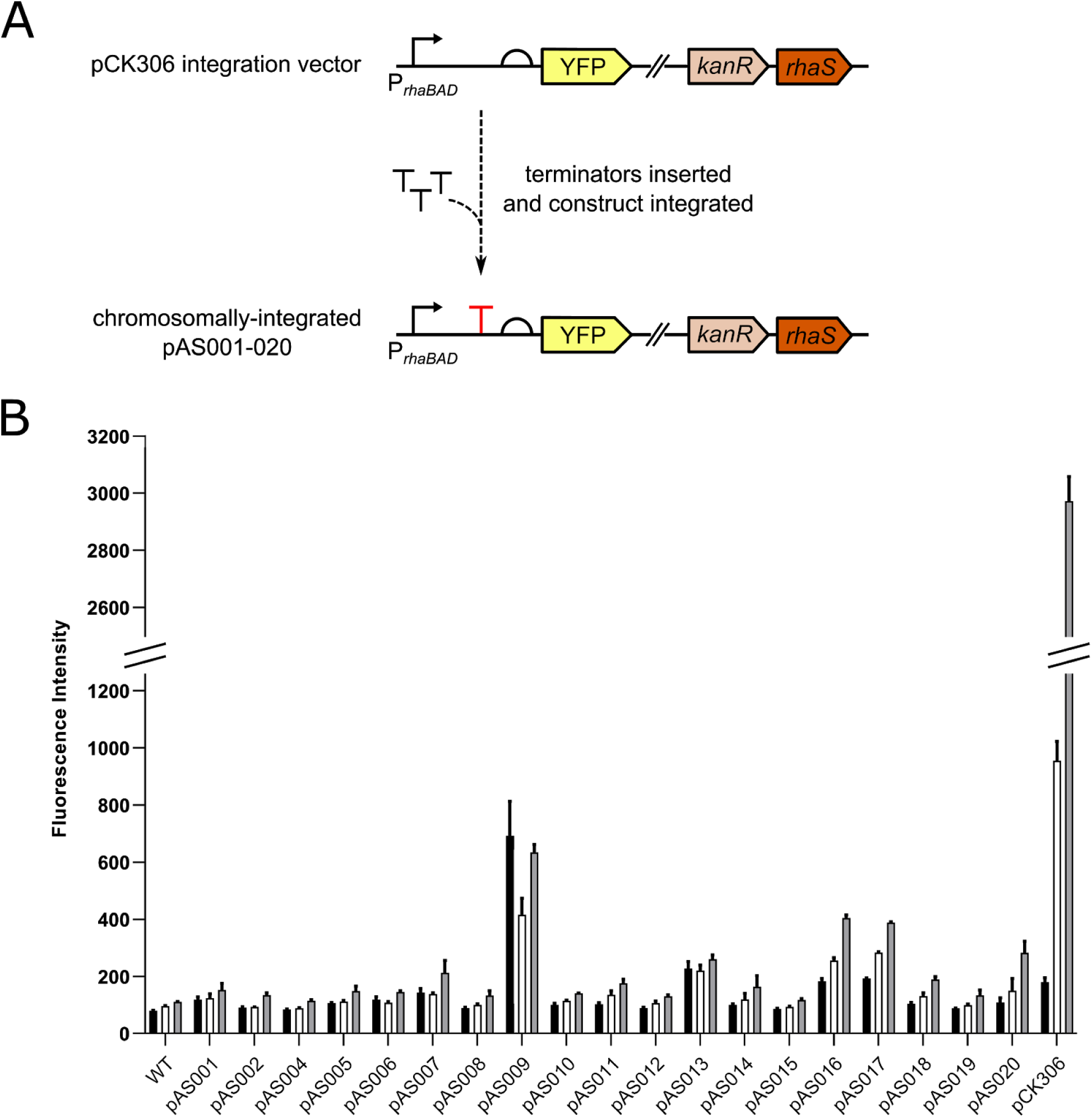
Screening of terminators in *Synechocystis*. (A) Overview showing insertion of terminators into plasmid pCK306 between the *rhaBAD* promoter and the RBS of the YFP-encoding gene. The resulting terminator constructs pAS001, pAS002, pAS004-020 were integrated into the *Synechocystis* genome. (B) *Synechocystis* transformants containing the integrated terminator constructs were cultured in BG11 media supplemented with kanamycin and 0.6 mg/ml L-rhamnose in photoautotrophic conditions in constant light. Wild-type *Synechocystis* cells (WT) lacking YFP entirely and cells containing the YFP-encoding cassette of the parent plasmid pCK306 (no terminator between *rhaBAD* promoter and RBS of YFP-encoding gene) were used as controls. The fluorescence intensity (arbitrary units) of 10,000 cells from each culture was measured by flow cytometry at 0 h, 48 h and 96 h (black, white and grey bars respectively). Error bars shown are the standard deviation of the mean for three independent biological replicates.

Following characterisation in *E. coli*, the terminator-containing plasmids pAS001, pAS002, pAS004-pAS020 were integrated into the *Synechocystis* genome with full segregation confirmed by PCR. The resulting transformants were cultured photoautotrophically, at 30 °C under constant light in BG11 growth medium supplemented with kanamycin and 0.6 mg/ml L-rhamnose. YFP fluorescence was measured by flow cytometry after 0, 48 and 96 h (linear phase of growth, as shown in Figure S3) and compared to wild-type (no YFP) *Synechocystis* cells and pCK306 (parent plasmid with no terminator inserted) transformants as controls (Figure 1). All pAS001, pAS002, pAS004-pAS020 transformants showed lower fluorescence than pCK306 transformants at 48 h and 96 h (highly statistically-significant difference; *P* value < 0.0001), indicating YFP production is reduced by the presence of any of these terminators. No significant difference in fluorescence was observed between wild-type cells and fifteen of the terminator-plasmid transformants at 48 h, with pAS009, pAS013, pAS016 and pAS017 showing significant differences (*P* values of <0.0001 for all). At 96 h, no significant difference in fluorescence was observed between wild-type cells and twelve of the terminator-plasmid transformants, with pAS007, pAS009, pAS013, pAS016, pAS017, pAS018 and pAS020 transformants showing significant differences (respective *P* values of < 0.001, < 0.0001, < 0.0001, < 0.0001, < 0.0001, < 0.05 and < 0.0001). These data show that twelve of the nineteen terminators tested in *Synechocystis* are strong transcriptional terminators, as they lead to YFP levels similar to those found in cells lacking YFP altogether. This conclusion is further supported by analysis of each transformant over the three timepoints, where no significant change in fluorescence was observed across the entire time course for eleven of these terminators. A significant difference (*P* value <0.005) was only observed for pAS011 between 0 h and 96 h, but no significant difference between either 0 h and 48 h or between 48 h and 96 h.

### Strong terminators insulate expression constructs integrated in the *Synechocystis* chromosome from transcriptional read-through

Four of the eleven strong terminator sequences were used to attempt to insulate integrated expression constructs from chromosomal read-through. These were terminators ECK120034435, ECK120015170, ECK120010799 and the *E. coli ilvBN* terminator, as originally screened using plasmids pAS004, pAS012, pAS014 and pAS019, respectively. Each of these terminators was introduced immediately upstream of the *rhaBAD* promoter of pCK306 (Figure 2A), resulting in plasmids pCK351, pCK353, pCK354 and pCK355, respectively. The four constructs were integrated into the *Synechocystis* genome at the locus adjacent to *ndhB* and full segregation was confirmed by PCR. *Synechocystis* transformants containing integrated pCK351, pCK353, pCK354 and pCK355 constructs were cultured at 30 °C under constant light in BG11 growth medium supplemented with kanamycin, with glucose (mixotrophic) or without glucose (photoautotrophic) and with or without 0.6 mg/ml L-rhamnose. YFP fluorescence was measured by flow cytometry after 0 h, 96 h and 192 h (linear phase of growth, as shown in Figure S4) and compared to wild-type (no YFP) *Synechocystis* cells and pCK306 (parent plasmid with no terminator inserted) transformants as controls. In the absence of inducer, the introduction of each of the four terminators reduced the YFP fluorescence of these cells by 1.7 - 2.1 times compared to pCK306 fluorescence (highly statistically-significant differences; *P* values of < 0.0001) in both photoautotrophic and mixotrophic growth conditions when measured at 96 h (Figure 2B and C). This resulted in very low fluorescence levels that were indistinguishable from wild-type cells lacking YFP entirely (no statistically-significant difference). The same pattern of results was observed at the later 192 h timepoint (Figure S2B and C). These data indicate that these four terminators are indeed effective at preventing transcriptional read-through from a chromosomal promoter and basal transcription has been reduced to undetectable levels. The four *Synechocystis* transformants containing YFP expression constructs insulated by upstream terminators were next tested to ensure inducible YFP production was still possible. Under photoautotrophic growth conditions, no significant difference in successfully-induced fluorescence was observed between each of the four transformants and the parent control pCK306 transformant at either 96 h (Figure 2D) or 192 h (Figure S2D). Under mixotrophic growth conditions, no significant difference in successfully-induced fluorescence was observed at either 96 h or 192 h, when pCK351 and pCK353 transformants were compared to the control transformant pCK306 (Figure 2E). A lower level of YFP was observed in transformants containing pCK354 and pCK355 constructs at the 96 h timepoint compared with pCK306 transformants (highly statistically-significant difference *P* value <0.0001) (Figure 2E). This difference had disappeared however at the later 192 h timepoint (Figure S2E).

**Figure 2.**
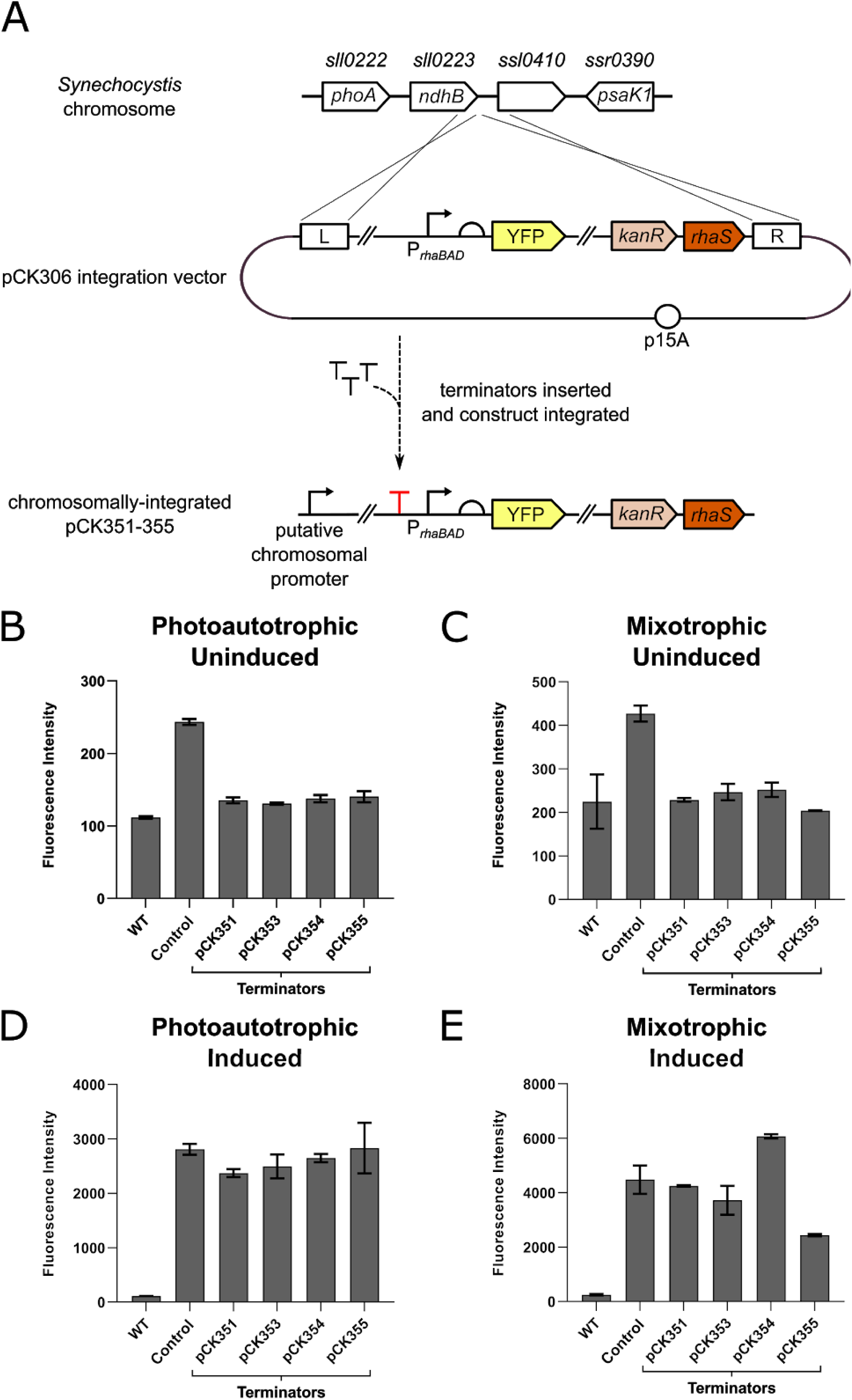
The effect of terminators inserted upstream of a rhamnose-inducible YFP expression construct inserted in the *Synechocystis* chromosome. (A) Detail showing the insertion of terminators into integration plasmid pCK306 upstream of the *rhaBAD* promoter. The resulting constructs pCK351, pCK353, pCK354 and pCK355 were integrated into the *Synechocystis* genome adjacent to the *ndhB* gene. (B) To test for transcriptional insulation from chromosomal promoters after integration, *Synechocystis* transformants containing the integrated terminator constructs were cultured in BG11 media supplemented with kanamycin and no L-rhamnose, in photoautotrophic conditions and constant light. Wild-type *Synechocystis* cells (WT) lacking YFP entirely and cells containing pCK306 (no terminator inserted upstream of *rhaBAD* promoter) were used as controls. The fluorescence intensity of 10,000 cells from each culture was measured by flow cytometry at 96 h. (C) Equivalent experiment to (B) but strains cultured in BG11 supplemented with 5 mM D-glucose (mixotrophic growth). (D) The same strains of *Synechocystis* were cultured in BG11 media supplemented kanamycin and L-rhamnose to a final concentration of 0.6 mg/ml in photoautotrophic conditions and constant light. The fluorescence intensity (arbitrary units) of 10,000 cells measured after 96 h using flow cytometry. (E) Equivalent experiment to (D) but strains cultured in BG11 supplemented with 5 mM D-glucose (mixotrophic growth). Error bars shown represent the standard deviation of the mean of three independent biological replicates.

## Discussion

In this study, nineteen Rho-independent terminators were evaluated in *Synechocystis*. This is important, as although a small number of *E. coli*-derived terminator sequences have previously been used in genetic constructs in cyanobacteria [2,8,9], no characterisation of their efficacy had been performed until now. The subunits of RNA polymerase differ between *E. coli* and *Synechocystis* and it has been reported that differences in RNA polymerase can lead to different termination efficiencies of terminators in different organisms [11]. Therefore, it was not obvious that all strong *E. coli* terminators would work well in *Synechocystis*. During preparation of this manuscript the characterisation of seven putative terminators found in the *Synechocystis* genome was reported [12], with moderate termination efficiencies observed. In our study, non-native and synthetic sequences were included and twelve strong Rho-independent terminators identified.

We demonstrated that four of the strongest terminators allow complete insulation of a chromosomally-integrated expression construct from transcriptional read-through resulting from genomic promoter(s) neighbouring the site of integration, presumably promoters of one or more of the sll0222 (*phoA*), sll0223 (*ndhB*) or ssl0410 genes [13]. In particular, we used strong terminators to improve the cyanobacterial rhamnose-inducible *rhaBAD* expression system previously reported [3,14], reducing basal transcription to undetectable levels. Importantly, this terminator insulation did not affect the inducible control of the expression system. The terminators characterised in this work are a valuable addition to the toolbox for engineering of cyanobacteria for fundamental and biotechnology purposes, allowing the insulation and independent control of individual expression units encoding separate components of a pathway or system. We recently included terminators when constructing metabolic pathways for cyanobacteria using the organism-independent Start-Stop Assembly system [15], as will be described in a forthcoming publication (Taylor and Heap, unpublished results). The improved and fully-insulated rhamnose-inducible *rhaBAD* expression system (plasmids pCK351, pCK353, pCK354 and pCK355) represents the best combination of high-level induced expression and undetectable levels of basal (non-induced) expression reported for any inducible system in cyanobacteria to date.

## Materials and Methods

### Bacterial strains and Growth Conditions

*E. coli* strain DH10β was used for all plasmid construction and propagation and the wild-type K12 strain MG1655 used for terminator-verification assays. *Synechocystis* sp. PCC 6803 WT-G (the glucose-tolerant derivative of the wild type, originally a kind gift from the laboratory of Peter Nixon at Imperial College London) was used for all cyanobacterial experiments. *E. coli* was cultured in LB media at 37 °C with shaking at 240 rpm and *Synechocystis* was cultured in TES-buffered (pH 8.2) BG11 media [16] with 5 mM glucose (mixotrophic growth) or without glucose (photoautotrophic growth) at 30 °C with shaking at 150 rpm, supplemented with 30 μg ml^-1^ kanamycin where required. *Synechocystis* was grown in constant white light at 50 μmol m^−2^ s^−1^.

### Plasmid Construction

A table of all plasmids and oligonucleotides (Table S2) is provided in the Supplementary Information. All plasmid construction was carried out using standard molecular cloning methods. Full details are provided in the Supplementary Information.

### Strain Construction

*Synechocystis* transformants were constructed as previously described [3] and full segregation confirmed by PCR.

### Assays

*Synechocystis* transformants were cultured in the presence of 0.6 mg/ml L-rhamnose, with the YFP fluorescence of samples at various time points (indicated in text) assayed as previously described [3] by flow cytometry using an Attune NxT Flow Cytometer (ThermoFisher). Cells were gated using forward and side scatter, and YFP fluorescence (excitation and emission wavelengths: 488 nm and 525 nm [with 20 nm bandwidth] respectively) was measured. Histograms of fluorescence intensity were plotted, and mean statistics extracted. *E. coli* transformants were cultured in LB media with or without 0.6 mg/ml L-rhamnose for 6 h and YFP fluorescence assayed as previously described [14] by flow cytometry. Error bars shown represent the standard deviation of three independent biological repeats. Statistical significance was determined using a one-way ANOVA, followed by a Tukey’s multiple comparison test assuming unequal variance.

## Supplementary Materials

Supplementary results, figures, tables and materials and methods available online.

## Acknowledgments

This work was supported by the Biotechnology and Biological Sciences Research Council [BB/M011321/1 to JTH].

## Author Contributions

CK and JH designed the study; all authors contributed to experimental design; CK, AS and GT performed experiments; CK, GT and JH prepared the manuscript with input from AS and LD.

## Conflicts of Interest

The authors declare no competing financial interest.

