## Supplementary Information for "Transcriptional terminators allow leak-free chromosomal integration of genetic constructs in cyanobacteria"

#### Supplementary Results

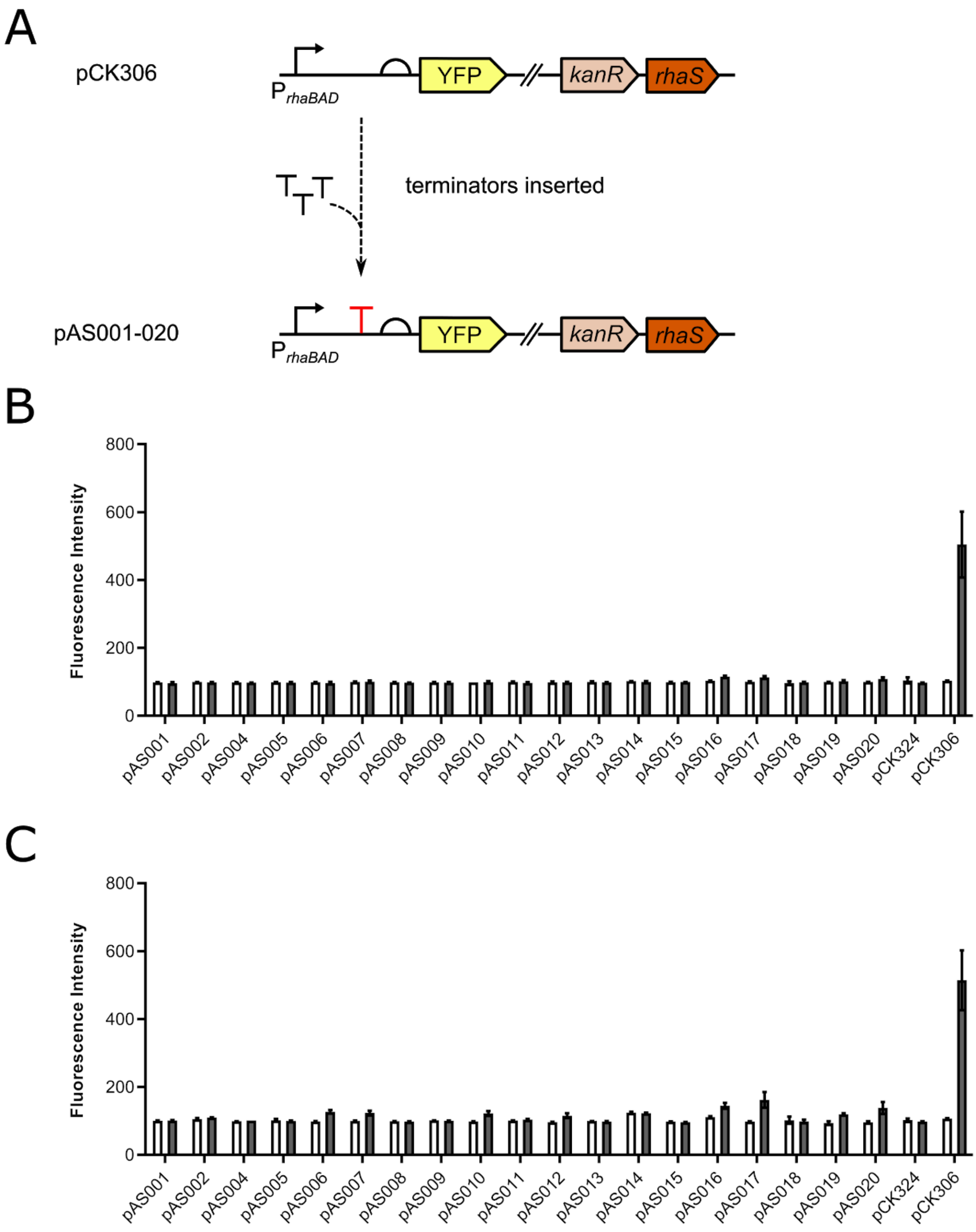
A
pCK306
P rhaBAD
YFP
kanR
rhaS
terminators inserted
pAS001-020
B
Fluorescence Intensity
800
600
400
200
0
pAS001
pAS002
pAS004
pAS005
pAS006
pAS007
pAS008
pAS009
pAS010
pAS011
pAS012
pAS013
pAS014
pAS015
pAS016
pAS017
pAS018
pAS019
pAS020
pCK324
pCK306
C
Fluorescence Intensity
800
600
400
200
0
pAS001
pAS002
pAS004
pAS005
pAS006
pAS007
pAS008
pAS009
pAS010
pAS011
pAS012
pAS013
pAS014
pAS015
pAS016
pAS017
pAS018
pAS019
pAS020
pCK324
pCK306

**Figure S1. Efficiency of terminators in *E. coli* strains DH5α and MG1655.** *E. coli* strains (A) DH10β and (B) MG1655 containing plasmids pAS001-002, pAS004-020 (terminator between *rhaBAD* promoter and RBS of YFP-encoding gene) were cultured in LB media supplemented with kanamycin and 0 mg/ml L-rhamnose (white bars) or 0.6 mg/ml L-rhamnose (black bars). Cells containing pCK324 (lacking *rhaBAD* promoter and therefore no YFP) and pCK306 (the parental vector with no terminator and therefore fully inducible with L-rhamnose) were used as controls. The fluorescence intensity of 10,000 cells (arbitrary units) from each culture was measured by flow cytometry after 6 h. Error bars shown are the standard deviation of the mean for three independent biological replicates.

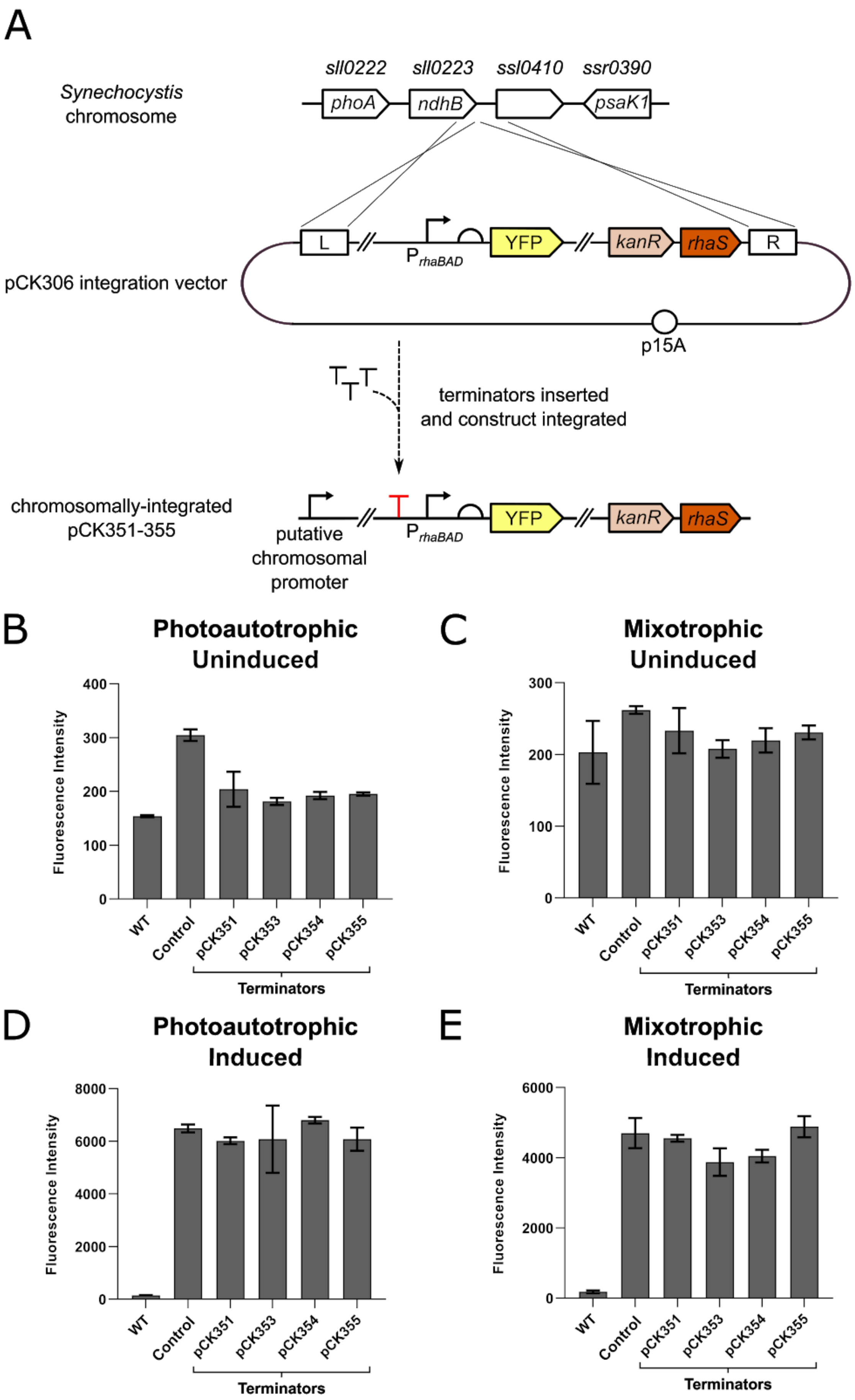
A
sll0222
sll0223
ssl0410
ssr0390
Synechocystis chromosome
phoA
ndhB
psaK1
pCK306 integration vector
L
P rhaBAD
YFP
kanR
rhaS
R
p15A
terminators inserted and construct integrated
chromosomally-integrated pCK351-355
putative chromosomal promoter
B
Photoautotrophic Uninduced
C
Mixotrophic Uninduced
D
Photoautotrophic Induced
E
Mixotrophic Induced
Fluorescence Intensity
WT
Control
pCK351
pCK353
pCK354
pCK355
Terminators

**Figure S2. The effect of terminator insertion upstream of chromosomally-integrated DNA on transcriptional read-through from chromosomal promoters.** (A) Detail showing the insertion of terminators into integration plasmid pCK306 upstream of the *rhaBAD* promoter. The resulting constructs pCK351, pCK353, pCK354 and pCK355 were integrated into the *Synechocystis* genome adjacent to the *ndhB* gene. (B) To test for transcriptional insulation from chromosomal promoters after integration, *Synechocystis* cells containing either pCK351, 353, 354 or 355 (each with one of four terminators inserted upstream of *rhaBAD* promoter) were cultured in BG11 media supplemented with kanamycin and no L-rhamnose, in photoautotrophic conditions and constant light. The fluorescence intensity of 10,000 cells measured after 192 h using flow cytometry and compared to wild-type and *Synechocystis* cells lacking YFP entirely and cells containing pCK306 (no terminator, *rhaBAD* promoter, YFP). (C) Equivalent experiment to (B) but strains cultured in BG11 supplemented with 5 mM D-glucose (mixotrophic growth). (D) The same strains of *Synechocystis* were cultured in BG11 media supplemented kanamycin and L-rhamnose to a final concentration of 0.6 mg/ml in photoautotrophic conditions and constant light. The fluorescence intensity of 10,000 cells (arbitrary units) measured after 192 h using flow cytometry and compared to wild-type and *Synechocystis* cells (lacking YFP entirely) and cells containing pCK306 (no terminator, *rhaBAD* promoter, YFP). (E) Equivalent experiment to (D) but strains cultured in BG11 supplemented with 5 mM D-glucose (mixotrophic growth). Error bars shown are the standard deviation of the mean for three independent biological replicates.

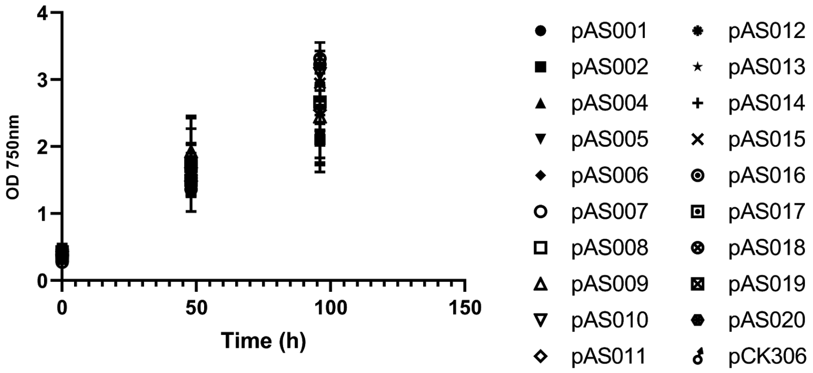

**Figure S3. Growth of *Synechocystis* cells transformed with terminator plasmids, pAS001-pAS020.** *Synechocystis* cells containing integrated terminator constructs from one of pAS001-002, pAS004-020 (terminator between *rhaBAD* promoter and RBS of YFP-encoding gene) or pCK306 (control, no terminator) were cultured in BG11 media supplemented with kanamycin and 0.6 mg/ml L-rhamnose in photoautotrophic conditions and constant light; and the optical density at 750 nm monitored over time. Error bars represent the standard deviation of the mean for three independent biological replicates.

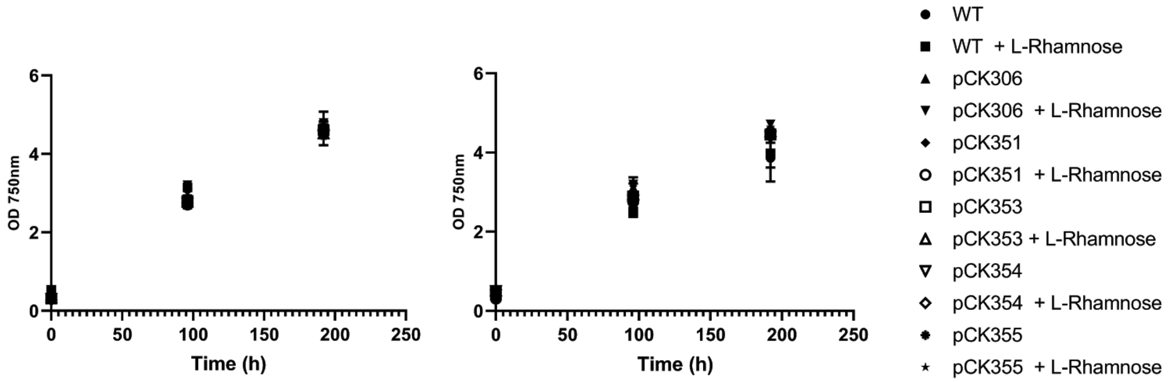

**Figure S4. Growth of *Synechocystis* cells transformed with insulated plasmids pCK351, pCK353, pCK354 or pCK355.** Wild-type (WT) *Synechocystis* cells, or cells containing integrated terminator constructs from one of pCK351, pCK353-355 plasmids (each with one of four Rho-independent terminators inserted upstream of the *rhaBAD* promoter) or pCK306 (control, no terminator) were cultured in BG11 media supplemented with kanamycin and 0 or 0.6 mg/ml L-rhamnose in photoautotrophic conditions and constant light; and the optical density at 750 nm monitored over time. Error bars represent the standard deviation of the mean for three independent biological replicates.

| Terminator | Screening plasmid | Length (bp) | Sequence (5'-3') | ΔG (kcal/mol) | Origin | References |
|---|---|---|---|---|---|---|
| ECK120029600 | pAS001 | 90 | TTCAGCCAAAAAACTTAAGACCGCCGGTCTTGTCCACTACCTTGCAGTAATGCGGTGGACAGGATCGGCGGTTTTCTTTTCTCTTCTCAA | -42.00 | *E. coli* K12 | Chen *et al*., 2013 |
| ECK120033737; *thrL* attenuator | pAS002 | 57 | GGAAACACAGAAAAAAGCCCGCACCTGACAGTGCGGGCTTTTTTTTTCGACCAAAGG | -25.00 | *E. coli* K12 | Chen *et al*., 2013; Jeng *et al*., 1990 |
| ECK120034435 | pAS004 | 57 | CTCGGTACCAAATTCCAGAAAAGAGACGCTGAAAAGCGTCTTTTTTCGTTTTGGTCC | -27.90 | *E. coli* K12 | Chen *et al*., 2013 |
| L3S2P21 | pAS005 | 61 | CTCGGTACCAAATTCCAGAAAAGAGGCCTCCCGAAAGGGGGGCCTTTTTTCGTTTTGGTCC | -37.90 | Synthetic | Chen *et al*., 2013 |
| L3S2P56 | pAS006 | 57 | CTCGGTACCAAATTTTCGAAAAAAGACGCTGAAAAGCGTCTTTTTTCGTTTTGGTCC | -28.80 | Synthetic | Chen *et al*., 2013 |
| L3S2P51 | pAS007 | 57 | CTCGGTACCAAAAAAAAAAAAAAAAGACGCTGAAAAGCGTCTTTTTTCGTTTTGGTCC | -24.90 | Synthetic | Chen *et al*., 2013 |
| L3S1P56 | pAS008 | 52 | TTTTCGAAAAAAGGCCTCCCAAATCGGGGGGCCTTTTTTATTGATAACAAAA | -23.40 | Synthetic | Chen *et al*., 2013 |

| | | | | | | |
|---|---|---|---|---|---|---|
| Bba_B0015; *rrnB* terminator | pAS009 | 130 | CCAGGCATCAAATAAAACGAAAGGCTCAG TCGAAAGACTGGGCCTTTCGTTTTATCTG TTGTTTGTCGGTGAACGCTCTCTACTAGA GTCACACTGGCTCACCTTCGGGTGGGCC TTTCTGCGTTTATA | -72.10 | *E. coli* K12 | Huang *et al*., 2010 |
| ECK120035133 | pAS010 | 43 | ACTGATTTTTAAGGCGACTGATGAGTCGC CTTTTTTTTGTCT | -15.40 | *E. coli* K12 | Chen *et al*., 2013 |
| ECK120017009 | pAS011 | 44 | GATCTAACTAAAAAGGCCGCTCTGCGGC CTTTTTTCTTTTCACT | -16.20 | *E. coli* K12 | Chen *et al*., 2013 |
| ECK120015170 | pAS012 | 47 | ACAATTTTCGAAAAAACCCGCTTCGGCGG GTTTTTTTATAGCTAAAA | -20.10 | *E. coli* K12 | Chen *et al*., 2013 |
| ECK120033736 | pAS013 | 53 | AACGCATGAGAAAGCCCCCGGAAGATCA CCTTCCGGGGGCTTTTTTATTGCGC | -37.80 | *E. coli* K12 | Chen *et al*., 2013 |
| ECK120010799 | pAS014 | 60 | GTTATGAGTCAGGAAAAAAGGCGACAGA GTAATCTGTCGCCTTTTTTCTTTGCTTGCT TT | -33.60 | *E. coli* K12 | Chen *et al*., 2013 |
| BBa_B0010; *rrnB* terminator | pAS015 | 80 | CCAGGCATCAAATAAAACGAAAGGCTCAG TCGAAAGACTGGGCCTTTCGTTTTATCTG TTGTTTGTCGGTGAACGCTCTC | -42.90 | *E. coli* K12 | Geerts *et al*., 1995 |
| Ω*groEL* | pAS016 | 89 | GGTTTAGTAGACCGACTACCACTTTTCTC ATAAAATCCCAGGGAGGTTTCGGCCTCCC TTTTTTTCACTTGCTAAGCTCTCTTTCGTT T | -20.80 | *Synechocystis* sp. PCC 6803 | Jacobsen and Frigaard, 2014 |

| | | | | | | |
|---|---|---|---|---|---|---|
| T21 | pAS017 | 74 | ATTGAGCAAGTAGCAACACTATTCGCATA AGCTGCCGTTAGTGACTCTTAAGTTGCAA CGGTGGCTTTTTTTAT | -25.40 | Bacteriophage T21 | Cambray *et al*., 2013 |
| M13 Central | pAS018 | 85 | AAAGCAAGCTGATAAACCGATACAATTAA AGGCTCCTTTTGGAGCCTTTTTTTTTGGA GATTTTCAACATGAAAAAATTATTATT | -18.60 | Bacteriophage M13 | Cambray *et al*., 2013 |
| *ilvBN* terminator | pAS019 | 36 | AAGACCCCCGCACCGAAAGGTCCGGGGG TTTTTTTT | -24.40 | *E. coli* K12 | Chen *et al*., 2013; Cambray *et al*., 2013 |
| ECK120010793 | pAS020 | 34 | TACGTAAAAACCCGCTTCGGCGGGTTTTT ACTTT | -24.40 | *E. coli* K12 | Chen *et al*., 2013; Cambray *et al*., 2013 |

**Table S1. Terminators used in this study**

### Supplementary Materials and Methods

#### Plasmid Construction

A table of all plasmids and oligonucleotides (Table S2) is provided. Terminators were introduced as follows. Each terminator sequence was split in two at the hairpin-loop sequence and each part was included at the 5’ end of oligonucleotides that were then used to amplify pCK306. PCR fragments were then ligated by blunt-end ligation using the New England Biolabs site-directed mutagenesis kit and sequence verified.

| Name | Details |
|---|---|
| pCK306 | Medium copy plasmid (p15A), with 2054 nucleotides of homology to the *Synechocystis* sp. PCC 6803 chromosome, allowing integration of DNA after the first 34 nucleotides of ssl0410 (adjacent to *ndhB*), antibiotic resistance gene *kanR* encoding an aminoglycoside phosphotransferase, the *rhaBAD* promoter from *E. coli*, a synthetic RBS and eYFP, the *E. coli rhaS* RBS and gene inserted downstream of the *kanR* gene. [1] |
| pAS001-20 | Detailed in Table S1 |
| pCK351 | As pCK306 but with terminator ECK120034435 inserted upstream of *rhaBAD* promoter |
| pCK353 | As pCK306 but with terminator ECK120015170 inserted upstream of *rhaBAD* promoter |
| pCK354 | As pCK306 but with terminator ECK120010799 inserted upstream of *rhaBAD* promoter |
| pCK355 | As pCK306 but with the *ilvBN* terminator inserted upstream of *rhaBAD* promoter |
| oligoAS001 | ACTGCAAGGTAGTGGACAAGACCGGCGGTCTTAAGTTTTTGGCTGAATA CGACCAGTCTAAAAAG<br>Used in construction of pAS001 |
| oligoAS002 | AATGCGGTGGACAGGATCGGCGGTTTTCTTTTCTCTTCTCAAATGAATCG GGTAAGTTTATAATATAC<br>Used in construction of pAS001 |
| oligoAS003 | AGGTGCGGGCTTTTTTCTGTGTTTCCTACGACCAGTCTAAAAAG<br>Used in construction of pAS002 |
| oligoAS004 | GACAGTGCGGGCTTTTTTTTCGACCAAAGGATGAATCGGGTAAGTTTAT AATATAC<br>Used in construction of pAS002 |
| oligoAS007 | TCTGGAATTTGGTACCGAGTACGACCAGTCTAAAAAG |

Used in construction of pAS004

oligoAS008 AAAGAGACGCTGAAAAGCGTCTTTTTTCGTTTTGGTCCATGAATCGGGTA
AGTTTATAATATAC
Used in construction of pAS004

oligoAS009 TCGGGAGGCCTCTTTTCTGGAATTTGGTACCGAGTACGACCAGTCTAAAA
AG
Used in construction of pAS005

oligoAS010 AAGGGGGGCCTTTTTTCGTTTTGGTCCATGAATCGGGTAAGTTTATAATAT
AC
Used in construction of pAS005

oligoAS011 TCAGCGTCTTTTTTCGAAAATTTGGTACCGAGTACGACCAGTCTAAAAAG
Used in construction of pAS006

oligoAS012 AAAGCGTCTTTTTTCGTTTTGGTCCATGAATCGGGTAAGTTTATAATATAC
Used in construction of pAS006

oligoAS013 TCAGCGTCTTTTTTTTTTTTTTTGGTACCGAGTACGACCAGTCTAAAAAG
Used in construction of pAS007

oligoAS014 AAAGCGTCTTTTTTCGTTTTGGTCCATGAATCGGGTAAGTTTATAATATAC
Used in construction of pAS007

oligoAS015 TTTGGGAGGCCTTTTTTCGAAAATACGACCAGTCTAAAAAG
Used in construction of pAS008

oligoAS016 TCGGGGGGCCTTTTTTATTGATAACAAAAATGAATCGGGTAAGTTTATAAT
ATAC
Used in construction of pAS008

oligoAS017 TCGACTGAGCCTTTCGTTTTATTTGATGCCTGGTACGACCAGTCTAAAAAG
Used in construction of pAS009

oligoAS018 AAGACTGGGCCTTTCGTTTTATCTGTTGTTTGTCGGTGAACGCTCTCTACT
AGAGTCACACTGGCTCACCTTCGGGTGGGCCTTTCTGCGTTTATAATGAA
TCGGGTAAGTTTATAATATAC
Used in construction of pAS009

oligoAS019 TCAGTCGCCTTAAAAATCAGTTACGACCAGTCTAAAAAG
Used in construction of pAS010

oligoAS020 TGAGTCGCCTTTTTTTTGTCTATGAATCGGGTAAGTTTATAATATAC
Used in construction of pAS010

oligoAS021 AGCGGCCTTTTTAGTTAGATCTACGACCAGTCTAAAAAG
Used in construction of pAS011

oligoAS022 CTGCGGCCTTTTTTCTTTTCACTATGAATCGGGTAAGTTTATAATATAC
Used in construction of pAS011

oligoAS023 AAGCGGGTTTTTTCGAAAATTGTTACGACCAGTCTAAAAAG
Used in construction of pAS012

oligoAS024 CGGCGGGTTTTTTTATAGCTAAAAATGAATCGGGTAAGTTTATAATATAC
Used in construction of pAS012

oligoAS025 ATCTTCCGGGGGCTTTCTCATGCGTTTACGACCAGTCTAAAAAG
Used in construction of pAS013

oligoAS026 CACCTTCCGGGGGCTTTTTTATTGCGCATGAATCGGGTAAGTTTATAATAT
AC
Used in construction of pAS013

oligoAS027 ACTCTGTCGCCTTTTTTCCTGACTCATAACTACGACCAGTCTAAAAAG
Used in construction of pAS014

oligoAS028 AATCTGTCGCCTTTTTTCTTTGCTTGCTTTATGAATCGGGTAAGTTTATAAT
ATAC
Used in construction of pAS014

oligoAS029 TTCGACTGAGCCTTTCGTTTTATTTGATGCCTGGTACGACCAGTCTAAAAA
G
Used in construction of pAS015

oligoAS030 AGACTGGGCCTTTCGTTTTATCTGTTGTTTGTCGGTGAACGCTCTCATGAA
TCGGGTAAGTTTATAATATAC
Used in construction of pAS015

oligoAS031 AAACCTCCCTGGGATTTTATGAGAAAAGTGGTAGTCGGTCTACTAAACCT
ACGACCAGTCTAAAAAG
Used in construction of pAS016

oligoAS032 CGGCCTCCCTTTTTTTCACTTGCTAAGCTCTCTTTCGTTTATGAATCGGGT
AAGTTTATAATATAC
Used in construction of pAS016

oligoAS033 AGAGTCACTAACGGCAGCTTATGCGAATAGTGTTGCTACTTGCTCAATTA
CGACCAGTCTAAAAAG
Used in construction of pAS017

oligoAS034 TAAGTTGCAACGGTGGCTTTTTTTATATGAATCGGGTAAGTTTATAATATA
C
Used in construction of pAS017

oligoAS035 AAGGAGCCTTTAATTGTATCGGTTTATCAGCTTGCTTTTACGACCAGTCTA
AAAAG
Used in construction of pAS018

oligoAS036 TTGGAGCCTTTTTTTTTGGAGATTTTCAACATGAAAAAATTATTATTATGAA
TCGGGTAAGTTTATAATATAC
Used in construction of pAS018

oligoAS037 TCGGTGCGGGGGTCTTTACGACCAGTCTAAAAAG

| | |
|---|---|
| | Used in construction of pAS019 |
| oligoAS038 | AAGGTCCGGGGGTTTTTTTTATGAATCGGGTAAGTTTATAATATAC<br>Used in construction of pAS019 |
| oligoAS039 | AAGCGGGTTTTTACGTATACGACCAGTCTAAAAAG<br>Used in construction of pAS020 |
| oligoAS040 | CGGCGGGTTTTTACTTTATGAATCGGGTAAGTTTATAATATAC<br>Used in construction of pAS020 |

**Table S2. Plasmids and oligonucleotides used in this study**
